# Bacterial Major Vault Protein homologs shed new light on origins of the enigmatic organelle

**DOI:** 10.1101/872010

**Authors:** Tymofii Sokolskyi

## Abstract

Vaults are large cone-shaped and highly conservative ribonucleoprotein complexes present in the cells of most major eukaryote clades. However, despite their wide distribution, their functions and evolutionary dynamics still remain enigmatic. Several minor functions in modulating signaling cascades and multidrug resistance phenotypes were previously discovered for eukaryotic vaults, yet nothing is known about bacterial homologs of the major vault protein (MVP), a protein that comprises the entirety of vault external surface. Using gene and protein BLAST searches in NCBI and UniProt databases we identified a number of bacterial species in prokaryotic taxa Myxococcales, Cytophagales and Oscillatoriales with >50% identity to eukaryotic MVP sequences. Interestingly, all of these species are characterized by one common feature – gliding type of motility. Secondary structures of the identified proteins were predicted using RAPTORX web service and aligned via jFATCAT-flexible algorithm in the RCSB PDB Java Structure Alignment tool to elucidate structural identity. Coiled coil domain at the MVP C-terminus of all studied bacterial species resembled TolA protein of *Escherichia coli* by both structure and sequence. We also showed that MVP sequences from chemotrophic bacteria Myxococcales and Cytophagales contain a domain homologous to eukaryotic band-7 domain, unlike cyanobacterial and eukaryotic major vault proteins. As expected, maximum-likelihood phylogenetic trees for MVP sequences separate studied taxa into two clades – first clade contains Oscillatoriales (Cyanobacteria) and Eukaryotes and the second one contains chemotrophic bacteria. In addition, binding prediction via RAPTORX showed great multiplicity GMP and CMP nucleoside monophosphate binding pockets in Myxococcales and Cytophagales MVP, unlike eukaryotic and cyanobacterial proteins which had much lower affinity to these substrates.

Due to high similarity of eukaryotic and cyanobacterial MVP sequences and a pattern of its phylogenetic distribution, we can speculate that the most likely scenario for vault appearance in eukaryotes is horizontal gene transfer from cyanobacteria. Presence of GMP and CMP binding pockets in MVP could also point to a function in depleting cytosolic nucleotide concentration which would be beneficial, for instance, during a viral infection. Further research is necessary to uncover potential functions of this enigmatic protein in bacteria and to determine its evolutionary patterns. In addition, a correlation between MVP presence and gliding motility in bacteria could also lead to elucidating selective pressures on the early evolution of this protein. Unfortunately, this topic has been largely neglected in recent literature and it can lead us to a much better understanding of not only current physiological processes but also eukaryogenesis, and even broader – origins of cellular life.

## Introduction

Vaults are large 13 MDa ribonucleoprotein complexes present in cells of many Eukaryota species (Kedersha et al., 1991). They consist of 3 types of proteins – major vault protein (MVP), vault poly-ADP ribose polymerase (vPARP), telomerase-associated protein (TEP1) and vault RNA. TEP1 is also shared with the telomerase complex, part of a so-called TROVE module that is shared between some ribonucleoproteins and mediates RNA binding (Bateman & Kickhoefer, 2003). The outer surface of a vault particle consists of MVP monomers forming two connected dome-shaped structures (Mikyas et al., 2004). It has been shown that vault particles are also capable of opening, possibly to transport other particles inside them (Querol-Audi et al., 2009). Vaults seem to be very conservative structures present among various Metazoa, Fungi and Protozoa taxa (Kong et al., 1999), meaning they could perform or could have performed a globally significant function. Multiple specific functions of this structure known in Metazoa are implicated in signaling pathway regulation, multidrug resistance, immunity, etc. (Berger et al., 2009). Currently vaults are even tested as drug and probe delivery vectors (Benner et al., 2017). These functions include:

1. Nuclear import of phosphoinositide phosphatase PTEN.
2. Scaffold in EGFR/MAPK cascade.
3. Apoptotic suppression via interactions with COP1 protein.
4. Nuclear import of activated estrogen receptors.
5. Unidentified role in axonal transport, possibly RNA transport (Li et al., 1999).
6. Possible regulation of poly-ADP ribosylation to facilitate DNA reparation.
7. Unidentified role in sea urchin ontogenesis (Stewart et al., 2005).
8. Multidrug resistance in mammal cells (Izquierdo et al., 1996).
9. Possible role in Epstein-Barr virus immunity.

A hypothesis was proposed for vaults to be an efficient way of amino acid and nucleotide storage (Shaik, 2013). This could explain their weird phylogenetic distribution – mostly in taxa that lost essential amino acid biosynthesis pathways and their upregulation during pathogen invasion – they could act as nutrient sequesters from the cytoplasm (Shaik, 2013). However, if this explanation is viable, then many questions regarding vault composition arise – what is the purpose of TEP1 and vPARP presence in the complex, why vaults contain relatively few RNAs, why are they shaped like they are and specifically why are they hollow inside.

Vault’s outer surface completely consists of major vault protein monomers. Recently its homologs were identified in several bacterial taxa, namely in some representatives of Cyanobacteria, Deltaproteobacteria and Bacteriodetes (Shaik, 2013). However, until now they have not been investigated neither *in silico* nor *in vivo*. Analysis of structure and evolution of bacterial MVP-like proteins may shed new light on vault function and evolution.

## Materials and methods

BLAST searches for the Homo sapiens MVP in Uniprot KB Bacteria database uncovered various proteins, several of which, representing different bacterial clades were selected for further investigation: *Moorea producens* uncharacterized protein F4Y3B4_9CYAN, *Saprospira grandis* uncharacterized protein H6L4P8_SAPGL, *Microscilla marina* uncharacterized protein A1ZGE7_9BACT, and *Enhygromyxa salina* uncharacterized protein A0A0C2D5V5_9DELT. Subsequent searches for these proteins found similarities with TolA proteins for various Proteobacteria and Bacteroidetes, and band 7 (SPFH) domain-containing proteins.

Then chosen MVP-like sequences from the first BLAST iteration were aligned with various TolA protein sequences found in the UniProt database for the respective taxa using local pairwise alignment tool EMBOSS Water (default settings, Smith-Waterman algorithm; Rice et al., 2000). Most common region of similarity was identified – sequence of approximately 200-250 amino acids. Maximum-likelihood phylogenetic tree with 1000 bootstrap replications were constructed in MEGA 7.0, including some sequences of eukaryotic MVP (Tamura et al., 2007).

Secondary structure and binding prediction for isolated regions of some of the studied proteins was performed with the help of RAPTORX web service (Källberg et al., 2012). Acquired structure data in the PDB format was then subjected for pairwise structure alignment via jFATCAT-flexible algorithm in the RCSB PDB Java Structure Alignment tool to confirm structural identity (Prli et al., 2010; Ye & Godzik, 2003).

## Results

### Comparison to TolA

BLAST searches for these sequences uncovered various TolA proteins; highest identity of 36% had *Gilliamella* TolA A0A1B9K2Y3_9GAMM. Series of alignments between the chosen sequences and TolA and TolA-like proteins of different bacterial species (*Enhygromyxa, Myxococcus, Sandaracinus, Escherichia, Haemophilius, Methylophaga, Mangrovibacter, Gilliamella, Frischella, Galibacterium*) showed that the highest similarity is located in the coiled coil region of the selected proteins and domain II of TolA. Identity variation between 22% and 37% and alignment of TolA sequences not only with the coiled coil region, but also with adjacent fragments of MVP shoulder and N-terminal domain suggests that it is most likely a result of homologous relationship between TolA domain II and MVP.

Results of the MOTIF search service on Genome Net (Bioinformatics Center at Kyoto University, http://www.genome.jp/tools/motif/), Pfam database (Finn et al., 2013) support this hypothesis showing TolA domains with E-values up to 1e-07 when analyzing bacterial MVP-like proteins. Schematic structure comparisons of *Saprospira* MVP-like protein and *Escherichia* TolA are shown in **Fig.1, A**.

**Fig.1.**
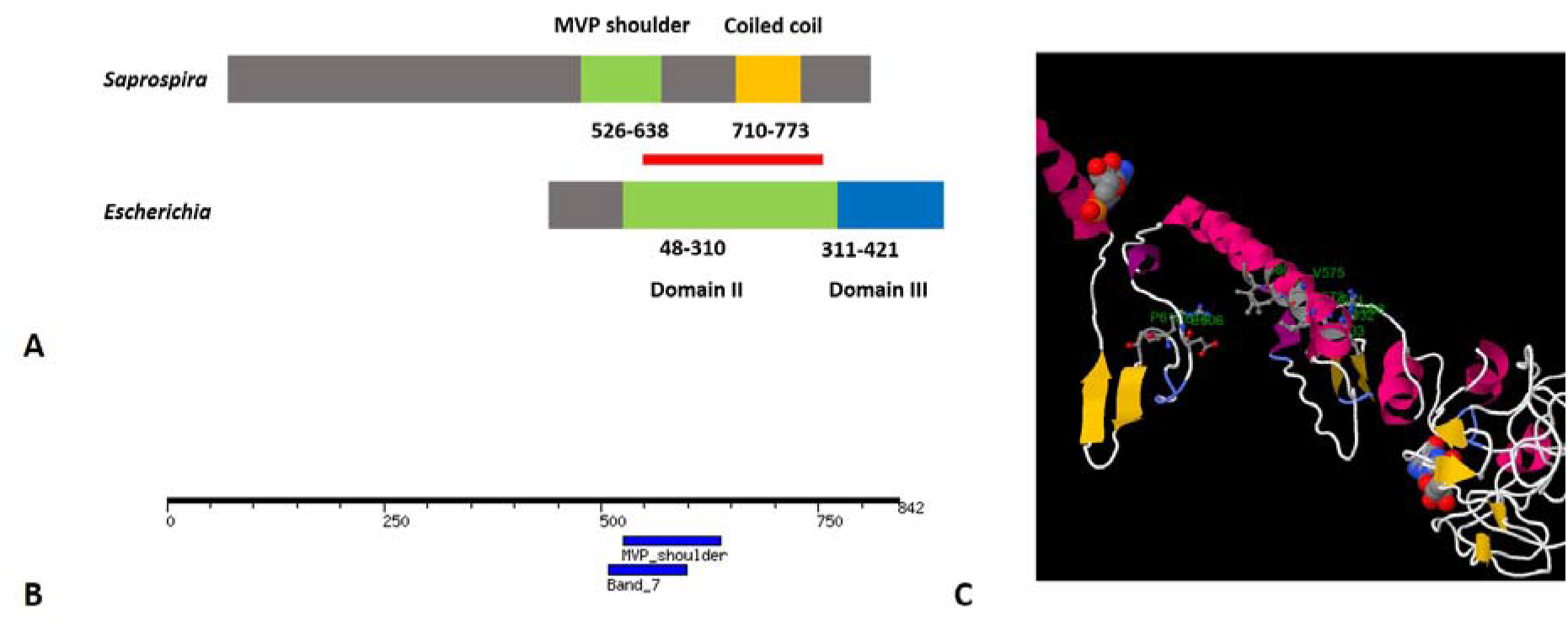
Structural features of MVP-like proteins. **A.** Schematic representation of *Saprospira grandis* uncharacterized protein (upper block) and *Escherichia coli* TolA protein (lower block) sequences. Aligned regions are marked with red line (portion of domain II from *E. coli* TolA and coiled coil with a small portion of MVP shoulder from *S. grandis* protein). Identity 27,6%, similarity 42,0%. **B.** Location of band 7 domain in *Saprospira* protein H6L4P8_SAPGL (result of MOTIF search). **C.** Graphic representation of a predicted GMP-binding site of *Saprospira* MVP-like protein.

### Structure prediction

Some of the most similar sequence fragments, corresponding to coiled coils of MVP-like proteins and domain II of TolA for some of the studied proteins were subjected to structure prediction via RAPTORX web service. Pairwise structural alignment was then conducted to support the idea of homology between major vault protein and TolA. Chosen sequences include: *Haemophilius* TolA TOLA_HAEIN (70-275); *Homo* Major Vault Protein MVP_HUMAN (729-805); *Moorea* MVP-like protein F4Y3B4_9CYAN (750-820); *Saprospira* uncharacterized protein H6L4P8_SAPGL (701-780). Acquired results, summarized in **table 1**, confirm the conclusion of evolutionary relationship between TolA and MVP-like proteins because of relatively low P-values and RMSD.

**Table 1.** Results of jFATCAT-flexible structure alignment of structures, explained in text. First number represents P-value, second – RMSD value.

|  | *Moorea* | *Homo* | *Haemophilus* | *Saprospira* |
| --- | --- | --- | --- | --- |
| *Moorea* |  |  |  |  |
| *Homo* | 1,79e-04; 3,17 |  |  |  |
| *Haemophilus* | 2,03e-05; 2,96 | 9,93e-04;<br>1,56 |  |  |
| *Saprospira* | 1,94e-08; 3,05 | 2,19e-06;<br>1,17 | 5,34e-06;<br>1,76 |  |

### Comparison with band 7 domains

Other than TolA, some BLAST searches for MVP shoulder-like domains in KB Bacteria database recovered more than 30% identity with band 7 proteins (*Halothermothrix* protein B8D0R4_HALOH showed the highest result of 33%). In addition, results of the MOTIF search service also identified SPFH / Band 7 domain in 509-598 region of *Saprospira* sequence H6L4P8_SAPGL with E-value=0.00039 (**Fig. 1, B**). Interestingly, same results were observed with other studied bacterial proteins, except cyanobacterial MVP-like proteins and eukaryotic MVP. This similarity was first reported by Daly et al., 2013.

Secondary structures for corresponding fragments of these two proteins along with *Pongo* MVP MVP_PONAB shoulder domain (519-647) and *Homo* stomatin STOM_HUMAN band 7 domain (56-227) were predicted and aligned using jFATCAT-flexible algorithm. Resulting RMSD and P-value are shown in **table 2**. P-values and RMSD are low enough to conclude likely homology. *Saprospira* MVP shoulder-like domain exhibits significantly higher structural similarity to band 7 domain of stomatin and *Halothermothrix* protein than to MVP shoulder of *Pongo* MVP.

**Table 2.** Results of jFATCAT-flexible alignment of structures, explained in text. First number represents P-value, second – RMSD value.

|  | *Halothermothrix* | *Homo* | *Pongo* | *Saprospira* |
| --- | --- | --- | --- | --- |
| *Halothermothrix* |  |  |  |  |
| *Homo* | 1,18e-12; 1,55 |  |  |  |
| *Pongo* | 1,57e-01; 3,01 | 1,55e-01; 3,03 |  |  |
| *Saprospira* | 1,18e-14; 1,36 | 6,38e-10; 1,82 | 2,68e-01; 3,26 |  |

### Binding prediction

Binding prediction via RAPTOR-X server was conducted for *Saprospira*, *Enhygromyxa*, *Moorea* and *Homo* MVP-like proteins. *Saprospira* protein showed high-multiplicity binding with GMP and CMP. Multiplicity represents the frequency with which the selected pocket was found in a set of ligand-binding protein structures – generally, if the value is above 40 the pocket is highly likely true. Multiplicity for the first GMP-binding site is 102, for the second – 35 (shown on **Fig. 1, C**), for CMP-binding site – 68. For *Enhyngromyxa* it is 102 and 28 for GMP and 72 for CMP. For *Moorea* protein it is only 46 and 12 for GMP and 28 for CMP; *Homo* MVP shows 37 and 14 for 2 possible GMP-binding sites, 38 for UMP and none for CMP. All of the possible nucleotide binding sites are located in 500-700 sequence, that corresponds to the part that aligns with MVP shoulder/band 7 domain. TolA proteins were also subjected for similar analysis. Neither *Haemophilius* nor *Gilliamella* TolA show any nucleotide binding *in silico*.

## Discussion

### Origin of vaults

We found 2 likely homologies for bacterial MVP-like proteins: TolA and band 7 domain proteins. A maximum-likelihood phylogenetic tree was constructed for complete protein sequences of various bacterial MVP-like proteins (**Fig. 2**). Two macroclades can be defined: first one contains Bacteroidetes and Deltaproteobacteria, the second one – eukaryotic MVP and cyanobacterial MVP-like proteins. The main structural difference between them is presence of band 7 domain in the sequences of all first clade taxa, and its absence in all second clade taxa. *Hyalangium* protein A0A085WQF1_9DELT is the only found non-cyanobacterial MVP-like protein lacking band 7 domain, no similar deltaproteobacterial sequences were found. Band 7 domains are generally known for association with lipid membranes (Tavernarakis et al., 1999).

**Fig. 2.**
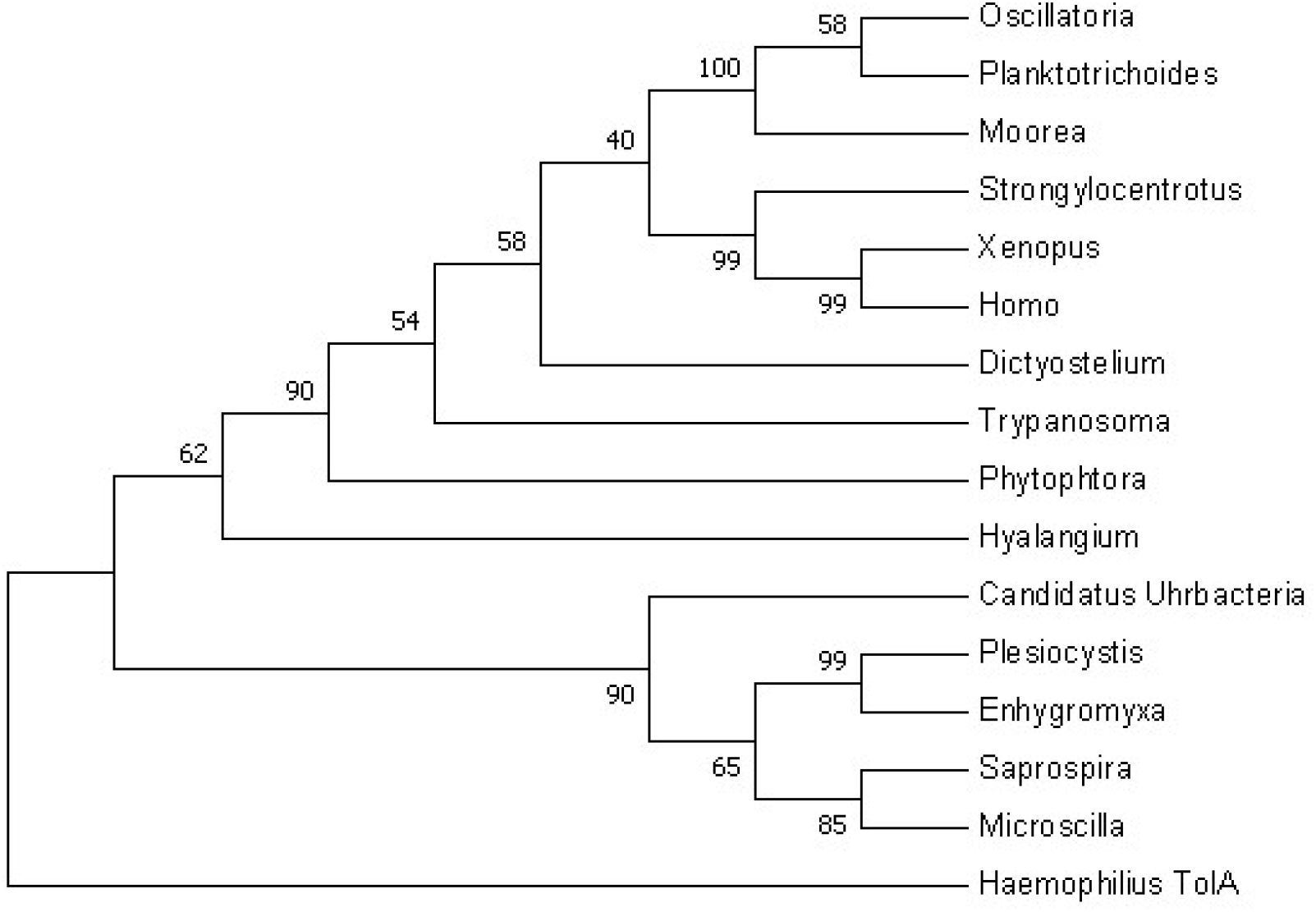
Maximum-likelihood phylogenetic tree derived for various bacterial MVP-like proteins. *Haemophilius* TolA sequence was used as an outgroup.

One of these bacteria, specifically *Saprospira grandis*, displays an unusual characteristic – rod-shaped protein structures in the cytoplasm, called rhaphidosomes (Saw et al., 2012). Their functions are currently unknown. One of the rhaphidosome proteins, SGRA_0791, was shown to contain Band 7-like domain (Saw et al., 2012). However, alignment of its 25-232 sequence that corresponds to Band 7 domain, according to Uniprot, with 500-600 sequence of *Saprospira* Band 7-like domain displayed only 20% identity, that is very low for such a short sequence, especially compared to >50% identities of MVP-like proteins between each other. At least for now there is no evidence of connection between rhaphidosomes and MVP-like proteins.

Major vault protein homologs were not found in archaea, that are closely related to eukaryotes. Therefore, MVP-like proteins were likely to appear in the bacterial lineage after Bacteria/Archaea divergence and then experience secondary loss in some lineages. Alternatively, they MVP-like proteins were present in the Last Universal Common Ancestor and then lost in Archaea and retained in Bacteria. However, if that is the case, then the loss should have occurred after the divergence of Eukarya and Archaea and then simultaneously happen in all contemporary archaeal clades which seems improbable. Therefore, we suggest that the most likely scenario of MVP appearance in Eukarya is lateral transfer from a bacterial lineage, most likely Cyanobacteria (Oscillatoriales), because they possess MVP-like protein with more than 50% identity to MVP. This event probably has happened before the divergence of major protist groups, animals, plants and fungi (Daly et al., 2013). This hypothesis is congruent with the phylogeny shown on **Fig. 2** that groups together cyanobacterial and eukaryotic MVP-like proteins.

Collected information suggests a probable evolutionary scenario for major vault protein, schematically depicted in **Fig. 3**. Pre-MVP appeared in the common ancestor of most bacterial phyla and contained band 7-like (SPFH) and TolAII-like domains (possibility of independent appearance of both domain types in many distantly related phyla is very low). Consequently, a duplication and a series of deletions occurred resulting in TolA protein, that lost band 7 domain in favor of a short transmembrane helix and acquired a variable N-terminal TolAIII domain. Some lineages, including Bacteroidetes, Cyanobacteria (at least Oscillatoriales) and Deltaproteobacteria retained the original pre-MVP. Ability to bind NMP appeared prior to Cyanobacteria/Bacteroidetes/Proteobacteria divergence, because it can be found in both *Moorea* and *Saprospira* proteins.

**Figure. 3.**
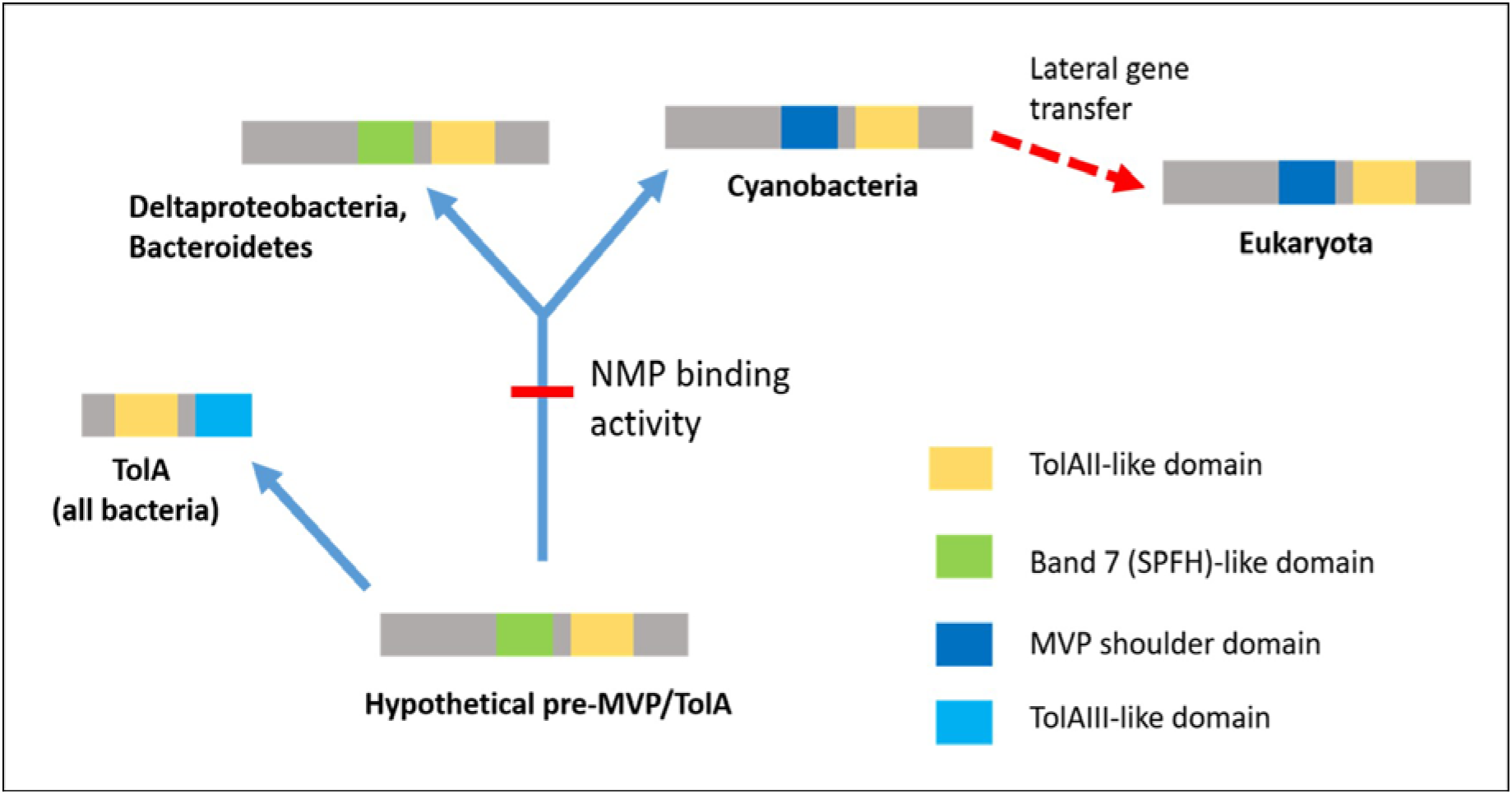
Most likely scenario of major vault protein evolution. Probably, NMP-binding appeared in Cyanobacteria and Deltaproteobacteria/Bacteroidetes MVP common ancestor, and then started waning in Cyanobacteria.

Cyanobacterial pre-MVP evolution made it move away from cell membrane transforming its band 7 domain into MVP shoulder. Cyanobacterial proteins could represent the first evolutionary steps towards the vault complex formation. Then, TolAII-like domain evolved to vault coiled coils that facilitate MVP aggregation into vaults, as demonstrated in modern eukaryotes (van Zon et al., 2002). Similar structure of *Hyalangium* protein could have been acquired convergently or due to a separate lateral gene transfer event. Then sequence of the resulting protein was laterally transmitted from cyanobacteria to early eukaryotes (most likely due to phagotrophic lifestyle of the latter ones (Shaik, 2013)) or due to an endosymbiotic event and was subsequently lost in various lineages possibly including plants and fungi (Daly et al., 2013).

### Future directions

All bacteria with MVP-like proteins identified here despite belonging to highly different taxa share a common characteristic: gliding motility (Engene et al., 2012; Iizuka et al., 2003 [1, 2]; Reichenbach, 2015). Specifically, these taxa include Deltaproteobacteria (only order Myxococcales), Bacteroidetes (specifically Cytophagales) and Cyanobacteria (only Oscillatoriales). This type of locomotion usually is prevalent among bacteria living in low-water content environments, including biofilms, mats, soils or hypersaline habitats, that is also true for the species studied here (Spormann, 1999). Interestingly, TolA protein that is partially homologous to bacterial MVP-like proteins is involved in flagellar locomotion (Cascales et al., 2001). Therefore MVP-like proteins could have played a role in the transition to gliding motility in selected taxa, mechanism of which is poorly studied (McBride, 2001).

Observation of MVP-like proteins NMP binding also supports the idea of vaults functioning as sequesters and regulators, as was proposed by Shaik, 2013. Bacterial MVP-like proteins could perform the same function vaults do, but without necessary aggregation into vaults. One of the interesting possible early functions of MVP-like proteins can be pathogen defense. By sequestering nucleotides and amino acids from the cytoplasm they can at least decrease the rate of bacteriophage reproduction. Nucleotides are required for bacteriophage replication, while they are not immediately necessary for bacterium during the infection process because no DNA replication is occurring (**Fig. 4**). It is possible that MVP demonstrates a lot weaker nucleotide binding than MVP-like proteins because of eukaryotic compartmentalization – deoxynucleotide concentration is a lot smaller in eukaryotic cytosol than in bacterial, meaning there is less sense in sequestering it during bacterial/viral infection. However, eukaryotes face wider systematic range of pathogens, meaning that a new way of spatial nutrient isolation should have been developed – that is aggregation into large hollow structures (vaults).

**Figure. 4.**
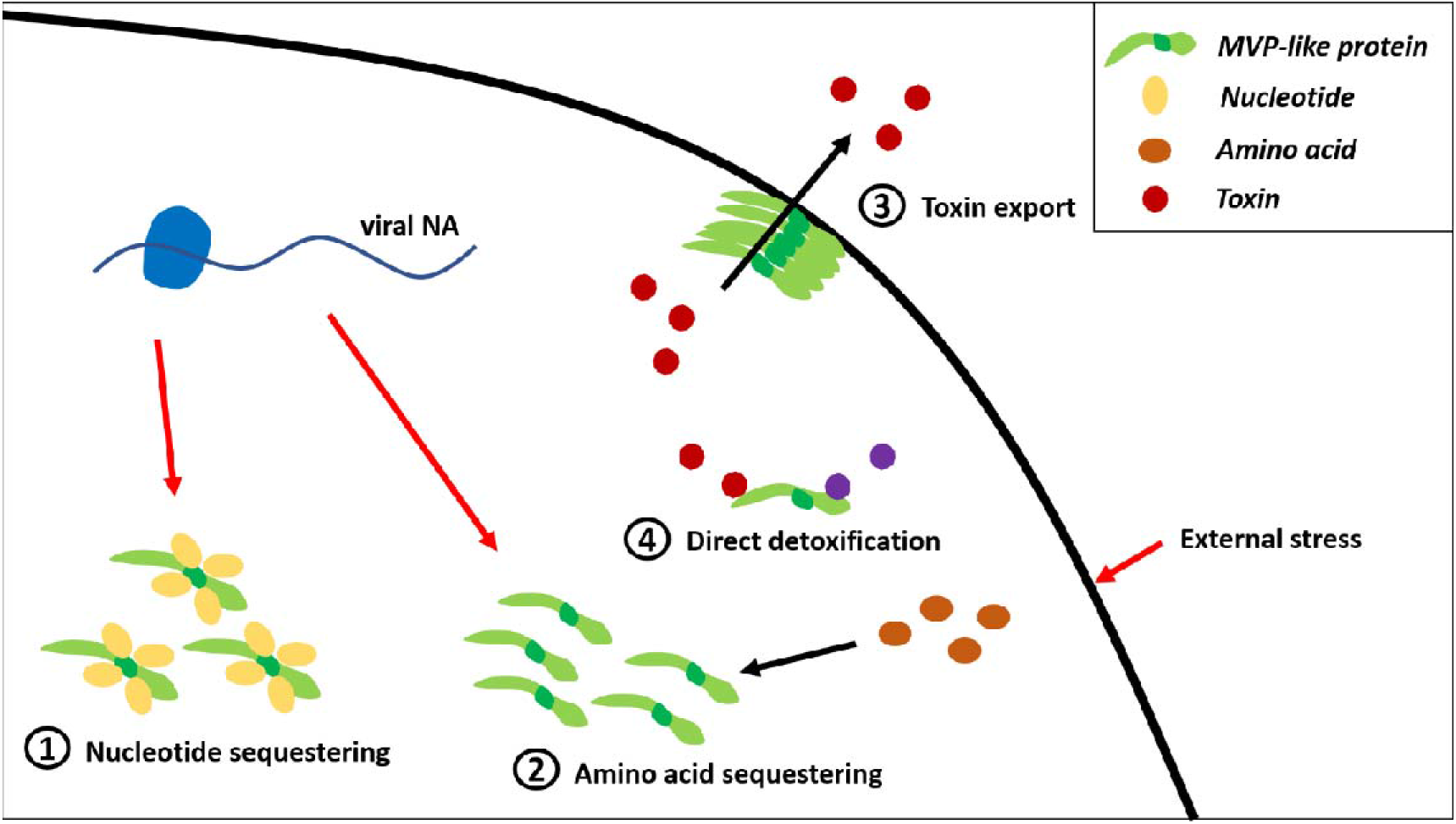
Possible functions of bacterial MVP-like proteins in cell protection. 1 – nucleotide sequestering from the cytosol to prevent viral replication; 2 – amino acid sequestering from the cytosol to affect viral protein synthesis or to conserve energy; 3 – export of toxic substances due to oligomerization through band 7 domains and membrane binding; 4 – direct detoxification.

Membrane-association via band 7 domains also could help them maintain this function. It has been shown that band 7 domain proteins can form oligomeric ring structures in mitochondria (Tatsuta et al., 2005) and cyanobacteria (Boehm et al., 2009) that are membrane-associated. This leads to an idea that MVP-like proteins could have also been associated with toxin export, similar to modern vaults involved in multidrug resistant phenotypes (Izquierdo et al., 1996). Alternatively, they could have also performed direct detoxification of substances such as toxic anions, as was demonstrated for modern MVP and tellurite anions (Suprenant et al., 2007). However, these hypotheses are purely speculative and require experimental verification.

MVP is one of the very few protein groups that are conserved among both Bacteria and Eukarya (Kedersha et al., 1990). It has been suggested that they were already present in last eukaryotic common ancestor, however their discovery in bacteria significantly expands their importance (Daly et al., 2013). In addition, the potential of cyanobacteria-eukaryote lateral gene transfer is intriguing and deserves further investigation. Unfortunately, the vault system has been largely neglected in the scientific community showing lack of publications even despite advances in molecular techniques over the past 35 years since the discovery of vaults (Kedersha & Rome, 1986). However, due to its conservativity and wide distribution, elucidating functions of vaults and MVP-like proteins in both prokaryotes and eukaryotes and their evolutionary patterns is necessary to understand eukaryogenesis and the broader picture of origins of cellular life. More *in silico*, *in vivo* and *in vitro* studies need to be done on this topic so that we can understand why these systems were retained by so many different taxa and how did they influence our evolution.

